# ZIP9 is a Druggable Determinant of Sex Differences in Melanoma

**DOI:** 10.1101/2020.03.12.989160

**Authors:** Cristina Aguirre-Portolés, Riley Payne, Aspen Trautz, J. Kevin Foskett, Christopher A. Natale, John T. Seykora, Todd W. Ridky

## Abstract

Melanoma and most other cancers occur more frequently, and have worse prognosis, in males compared with females. Though sex steroids are thought to be involved, classical androgen and estrogen receptors are not detectable in most melanomas. Here we show that testosterone promotes melanoma proliferation by activating ZIP9 (*SLC39A9*), a zinc transporter that is not intentionally targeted by available therapeutics, but is widely expressed in human melanoma. This testosterone activity requires zinc influx, MAPK activation and YAP1 nuclear translocation. We demonstrate that FDA approved inhibitors of the classical androgen receptor also inhibit ZIP9, and thereby antagonize the pro-tumorigenic effects of testosterone in melanoma. In male mice, androgen receptor inhibitors suppressed growth of ZIP9-expressing melanomas, but had no effect on isogenic melanomas lacking ZIP9, nor on melanomas in females. These data suggest that ZIP9 might be effectively targeted in melanoma and other cancers by repurposing androgen receptor inhibitors that are currently approved only for prostate cancer.

**Significance:** Melanoma outcomes are worse in males than in females. Some of this difference is driven by testosterone signaling through ZIP9, a nonclassical testosterone receptor. Drugs that target AR can be repurposed to block ZIP9, and inhibit melanoma in males.

## Introduction

Cancer incidence and mortality are higher in males than in females in the U.S. and worldwide^1,2^. In the U.S., males are 15% more likely to develop cancer, and 40% more likely to die from this disease than females^1^. These sex differences were recognized as early as 1949^3^, are observed in the majority of cancer types from non-reproductive tissues^1^, and remain even after controlling for known risk factors such as environmental and occupational exposures^4^. While recent advances in modern targeted and immune therapeutics have markedly improved survival for both female and male melanoma patients^5^, females still have more favorable outcomes^1,4^. Defining the mechanisms underlying the broad and persistent sex differences in cancer incidence and outcomes will address a major unresolved question in cancer pathobiology.

We previously showed that the female sex hormone estradiol inhibits melanoma proliferation and tumor growth *in vivo*. This effect is independent of biologic sex of the tumor, independent of classical estrogen receptors (ER), and results from activation of a nonclassical surface estrogen receptor on melanocytes and melanoma cells called the G Protein Estrogen Receptor (GPER)^6,7^. This work led us to consider whether male sex hormone signaling might also contribute to female vs. male differences in melanoma progression.

Testosterone is the most abundant androgen and circulates at much higher levels in males (630 ng/dl) than in females (32 ng/dl)^8^. In males, higher levels of circulating testosterone also correlate with increased melanoma incidence^9^. Testosterone promotes proliferation *in vitro* in non-gonadal cell types including adipocytes^10^, mouse skeletal muscle myoblasts^11^, glioblastoma-derived cells^12^, lung cancer cell lines^13^ and melanoma^14^. As the classical androgen receptor (AR) is not consistently detected in most of these tissues, the receptor(s) mediating the testosterone-dependent increased proliferation and the corresponding downstream mechanisms are not yet defined. Also unknown is whether this androgen activity observed *in vitro* is relevant to cancer progression *in vivo*.

Here we studied 98 human melanocytic lesions (nevus, primary and metastatic melanoma from both males and females) and did not detect AR in any of them. However, in nearly all samples, we readily detected ZIP9, a membrane localized zinc transporter recently discovered in Atlantic Croaker (fish) cells to be a activated by testosterone^15^.

Here, we use genetic and pharmacologic approaches to target ZIP9 in multiple melanoma models *in vitro* and *in vivo*. These functional studies define a previously unappreciated nonclassical testosterone signaling pathway through ZIP9, establish a novel mechanistic link between male androgens and melanoma pathobiology, and highlight a potential new therapeutic opportunity by repurposing currently available drugs.

## Materials and Methods

### Cell culture and proliferation assays

YUMM1.7, SH-4 and SK-MEL-2 cells were purchased from ATCC (YUMM1.7 ATCC^®^ CRL-3362™; SH-4 ATCC^®^ CRL-7724™; SK-MEL-2 ATCC^®^ HTB-68™) and cultured in DMEM (Mediatech, Manassas, VA, USA) with 5% FBS (Invitrogen, Carlsbad, CA, USA) and 1% Penicillin-Streptomycin (Thermo Fisher Scientific. #15140122). SK-MEL-3 cells were purchased from ATCC (ATCC^®^ HTB-69™ and cultured in McCoy’s 5A (Modified) Medium with 10% FBS (Invitrogen, Carlsbad, CA, USA) and 1% Penicillin-Streptomycin. SK-MEL-24 cells were purchased from ATCC (ATCC^®^ HTB-71™) and cultured in Eagle’s Minimum Essential Medium with 15% FBS and 1% Penicillin-Streptomycin. WM46 melanoma cells were a gift from Meenhard Herlyn (Wistar Institute, Philadelphia, PA, USA) and were cultured in TU2% media. Tumor cells were regularly tested using MycoAlert Mycoplasma Detection Kit from Lonza (Allendale, NJ, USA). Early passage cells were used after obtaining them directly from ATCC. For monitoring cell proliferation 10×10^5^ YUMM1.7 or 12×10^5^ WM46 cells were seed per well in 12-well Cell culture Plates. All the experiments performed in this work utilized charcoal stripped serum (Fetal Bovine Serum, charcoal stripped, USDA-approved regions, One Shot™ format. Catalogue number #A3382101; Thermofisher Scientific). Cells were treated every second day and manually counted in triplicate using a hemocytometer (Testosterone solution and Testosterone-CMO-BSA were purchased from Sigma-Aldrich, T5411-1ML and T3392-10MG respectively). All the experiments were performed in cell populations that were in culture during a maximum of 3 weeks (5 passages in average) since thaw from the corresponding stock.

### CRISPR-Cas9 mediated ablation of Slc39A9

We used lentiviral transduction to deliver dox-inducible Cas9 and gRNA targeting exon 1 of Slc39A9 in human WM46 and murine YUMM1.7 melanoma cells. Transduced cells were selected with puromycin, and single cells subsequently isolated, expanded and examined for ZIP9 protein expression, compared to clones isolated in parallel with no doxycycline treatment. The following gRNA sequences were used (5’-3’):

hZIP9_gRNA_Fw caccgTTGGTGGGATGTTACGTGGC

hZIP9_gRNA_Rv aaacGCCACGTAACATCCCACCAAC

hZIP9_gRNA_Fw caccgCGTGGCCGGAATCATTCC

hZIP9_gRNA_Rv aaacGGAATGATTCCGGCCACG

To map the targeted sequencing, the region surrounding the gRNA target sequence was amplified in both WM46 (282bp) and YUMM1.7 (259bp) isogenic clones. The following primers were used (5’-3’): hZIP9_CRISPRmut_Fw:

TAAGCAGAATTCATGGATGATTTCATCTCC

hZIP9_CRISPRmut_Rv:TAAGTAAGTCCAAGCTTCTGCTGCTTTGTCTGATGCA.

mZIP9_CRISPRmut_F:TAAGCAGAATTCATGGATGACTTTCTCTC.

mZIP9_CRISPRmut_Rv TAAGTAAGTCCAAGCTTGATATTTCTGCTGCTTTGT.

Once amplified, DNA fragments were cloned into pUC19 vector using EcoRI and HindDIII sites, and sequenced (Sanger) by the DNA Sequencing Facility (University of Pennsylvania). Sequences were analyzed using the free software CRISP-ID (V1.1) and ICE Analysis by Synthego were the knock-out scores were obtained.

### Zinc influx analysis

WM46 cells were loaded with 5 µM FluoZin™-3, AM cell permeant™ (Thermo Fisher Scientific, #F24195), for 20 minutes then incubated in 2 Ca Tyrode’s solution (in mM: 140 NaCl, 5 KCl, 1 MgCl_2_, 2 CaCl_2_, 10 glucose and 5 (Na) Pyruvate – pH 7.4) for 5-10 minutes at room temperature prior to imaging. Cells were imaged on a Nikon Ti microscope using a 20x/0.75 NA objective for fluorescence at 340 nm and 380nm excitation/515 nm emission (Ca^2+^-free Fura2 or FluoZin™-3) and 380 nm excitation/515 nm emission (Ca^2+^-free Fura2 FluoZin™-3). Coverslips were perfused at 1-3 mL/min following this protocol: Ca^2+^ Tyrode’s (0-30 secs); Ca^2+^ Tyrode’s + DMSO & 5 uM Zinc (30-90 secs); Ca^2+^ Tyrode’s + Testosterone & 5μM Zinc (90-360 secs); Ca^2+^ Tyrode’s + Testosterone + BIC & 5 uM Zn (360-470 secs); Ca^2+^ Tyrode’s washout (470-500 secs). 100 milliseconds exposure images for each wavelength were collected every 2 seconds. For analyzing the long-term consequences of testosterone treatment, cells were treated with the androgen for 96 hours and then loaded FluoZin™-3. Cells were incubated with FluoZin™-3 following manufacturer recommendation and exposed to 400nM Zn pyrithione or ZnCl_2_. Images were acquired with EVOS BLA. Fluorescence was quantitated using ImageJ (National Institutes of Health, Bethesda, MD, USA), and statistical analyses were performed using Graphpad Prism software.

### Reverse Phase Protein Array (RPPA)

We used the Functional Proteomics Core at MD Anderson (https://www.mdanderson.org/research/research-resources/core-facilities/functional-proteomics-rppa-core.html) to perform a 447-element Reverse Phase Protein Array (RPPA) analysis of human melanoma cells grown in medium with stripped serum and treated with testosterone (100nM) for 0, 30’, 60’ and 8 hours. Cells were tripsinized and washed with PBS. After centrifugation (5 minutes; 1200rpm), supernatant was discarded, and cells were frozen at -80°C prior to be sent to the Functional Proteomics Core at MD Anderson.

### Western blot, immunofluorescence and antibodies

Adherent cells were washed once with PBS and lysed with 8M urea containing 50mM NaCl and 50mM Tris-HCl, pH 8.3, 10mM dithiothreitol, 50mM iodoacetamide. Lysates were quantified (Bradford assay), normalized, reduced, and resolved by SDS gel electrophoresis on 4–15% Tris/Glycine gels (Bio-Rad, Hercules, CA, USA). Resolved protein was transferred to PVDF membranes (Millipore, Billerica, MA, USA) using a Semi-Dry Transfer Cell (Bio-Rad), blocked in 5% BSA in TBS-T and probed with primary antibodies recognizing β-actin (Cell Signaling Technology, #3700. Mouse. Lot:14 1:4000, Danvers, MA, USA), ZIP9 (Abcam, #137205, Rabbit mAb. Lot: GR3231323-8. 1:500), P-ERK (Cell Signaling Technology, Phospho-p44/42 MAPK (Erk1/2) (Thr202/Tyr204) (D13.14.4E) XP^®^ Rabbit mAb #4370. Lot. 24 1:1000), ERK (Cell Signaling Technology, p44/42 MAPK (Erk1/2) (137F5) Rabbit mAb. Lot: 28 #4695, 1:1000), Androgen Receptor [(D6F11) XP^®^ Rabbit mAb #5153. Lot: 7], Recombinant Anti-Androgen Receptor antibody [EPR1535(2)] (ab133273. Antibody already validated by the Human Protein Atlas), RSK1/RSK2/RSK3 [(32D7) Rabbit mAb #9355. Lot:3], Phospho-p90RSK [(Thr359/Ser363)Antibody #9344. Lot: 15]. After incubation with the appropriate secondary antibody [(Rabbit Anti-Mouse IgG H&L (Biotin) preabsoFS9rbed (ab7074); Anti-mouse IgG, HRP-linked Antibody #7076. 1:2000)] proteins were detected using ClarityTM Western ECL Substrate (Bio-Rad. #170-5060). All western blots were repeated at least 3 times. To monitor YAP1 nuclear translocation by western blot analysis, nuclear fractionation was performed using protein lysis buffer containing 10mM HEPES, 1mM KCl, 1.5mM MgCl2 and 10% glycerol (Buffer A). After washing the adherent cells with DPBS, samples were resuspended in Buffer A and incubated for 5 minutes on ice in the presence of 0.1% Triton-X-100. After centrifugation, the nuclear fraction remained as a pellet while the supernatant corresponding to the cytosolic fraction. Nuclear fraction was washed with Buffer A. After centrifugation, the nuclear fraction was resuspended in RIPA buffer and samples were boiled for 5 min prior to sample loading. The cytosolic fraction was centrifuged for 15 min at 13500 rpm. Only the supernatant was kept after centrifugation. Western blot was performed as described before and the following antibodies were used for protein detection: β-tubulin (Cell Signaling Technology, β-Tubulin (9F3) Rabbit mAb #2128. 1:1000), PARP (Cell Signaling Technology, Rabbit mAb #9542. 1:1000), YAP1 (Cell Signaling Technology, YAP (D8H1X) XP^®^ Rabbit mAb #14074. 1:1000). For immunofluorescence in adherent cells, samples were fixed with 4% paraformaldehyde (Affymetrix; #19943) for 7 min at room temperature and permeabilized with iced-cold methanol. After blocking with 10% FBS:0.03%Triton-X100, primary antibodies were incubated overnight in blocking solution (β-actin and YAP1 antibodies detailed above). After three washes in DPBS:0.03% Triton X-100) cells were incubated with secondary antibodies for 45 minutes at room temperature [(Goat anti-Mouse IgG (H+L) Highly Cross-Adsorbed Secondary Antibody, Alexa Fluor 594. Goat anti-Rabbit IgG (H+L) Highly Cross-Adsorbed Secondary Antibody, Alexa Fluor 488)]. Cells were rinsed with PBS-0.03%-Triton three times and coverslips were mounted with ProLong Gold antifade reagent with DAPI (#P36935; Thermo Fisher Scientific). Images were captured using a Leica DM IL microscope and registered using LAS software. For fluorescence intensity quantification ImageJ software (National Institutes of Health, Bethesda, MD, USA) was used, and statistical analyses were performed with GraphPad Prism software.

### Immunohistochemistry and quantification

FFPE tissue microarrays (ME1004h: Malignant melanoma, metastatic malignant melanoma and nevus tissue array) were obtained from US Biomax, Inc. (Derwood, MD). For the staining with anti-ZIP9 antibody [(SLC39A9 Antibody (PA5-52485), Thermofisher Scientific, Waltham, MA)], slides were deparaffinized and rehydrated following the standard immunohistochemistry protocol [(xylenes 5 minutes x 3, 100% alcohol (5 min. x 3), 95% alcohol (5 min.), 80% alcohol (5 min.), 70% alcohol (5 min.), and 50% alcohol (5 min.) and finished with distilled water)]. The antigen retrieval was done by loading the slides into a retriever (Electron Microscopy Sciences EMS) with R-Buffer A. After 20 minutes, samples were allowed to cool for 30 minutes inside the retriever and for 20 minutes at room temperature. Samples were washed twice with PBS and blocked with Dako Dual Endogenous Enzyme Block (Code S2003. Agilent Santa Clara, CA) for 20 minutes at R.T. Samples were washed twice with PBS and blocked for 20 minutes with 2 drops of Vector Avidin Block. After washing twice with PBS, slides were blocked for 20 minutes with two drops of Vector Biotin Block (Avidin/Biotin Blocking kit. SP-2001. Vector Laboratories, Inc. Burlingame, CA). Samples were washed twice with PBS with Protein Block Serum-Free Ready-To-Use for 30 minutes at R.T. (Code X0909. Agilent Santa Clara, CA). Primary antibody was prepared 1:500 in PBST (100μl per slide) and samples were incubated overnight at 4°C. Samples were washed three times with PBS and incubated with Biotinylated Secondary antibody (Vectastain Kit, Peroxidase Rabbit IgG, PK-4001) for one hour at R.T. After three washes with PBS, ABC reagent (prepared 30 minutes in advance) was added and samples were incubated for 30 minutes at R.T. Samples were washed twice with PBS and incubated for 3 minutes with ImmPACT^®^ DAB Substrate, Peroxidase (HRP) (SK-4105. Vector laboratories, Burlingame, CA). Tissues were counterstained with hematoxylin (30 seconds, R.T.) (GHS316. Sigma-Aldrich) dehydrated, and mounted with SecureMount (Fisher HealthCare™ PROTOCOL™ Mounting Media. #022-208. Fisher Scientific. Thermofisher Scientific). John T. Seykora M.D., Ph.D, performed scoring of the stained tissue microarray, and scoring index was determined by scoring the percentage of positive cells on a scale of 0 to 3 as well as the intensity of ZIP9 staining on a scale of 0 to 4 (1=1-25%, 2=26-50%, 3=51-75%, 4=76-100%).

The staining of the tissue microarrays (ME1004h) for AR detection was performed by University of Pennsylvania Pathology Clinical Service Center—Anatomic Pathology Division, using the highest grade, CLIA (Clinical Laboratory Improvement Amendments) certified and validated test available. Briefly, five-micron sections of formalin-fixed paraffin-embedded tissue were stained using antibody against Androgen Receptor [(Leica AR-318-L-CE, clone AR27 (clone AR27, 1:25)]. Staining was done on a Leica Bond-IIITM instrument using the Bond Polymer Refine Detection System (Leica Microsystems DS9800). Heat-induced epitope retrieval was done for 20 minutes with ER2 solution (Leica Microsystems AR9640). All the experiment was done at room temperature. Slides are washed three times between each step with bond wash buffer or water. The slides were reviewed and scored in blinded fashion by a board-certified U. Penn pathologist. Prostate tissue was used as the positive control.

### Immunocytochemistry

To detect ZIP9 protein in WM46 isogenic clones, the protocol described in ProSciΨ™ for immunocytochemistry. Briefly, cells were fixed in 4% PFA for 7 minutes. After two washes with PBS (5min), cells were permeabilized with PBS/0.1% Triton X-100 for 1 minute at R.T. Cells were washed twice with PBS on a shaker. Once treated with 1.5% H_2_O_2_/PBS solution for 15 minutes (R.T.), cells were washed again and blocked with 5%BSA for one hour at R.T. For primary antibody incubation, α-ZIP9 antibody [(SLC39A9 Antibody (PA5-52485), Thermofisher Scientific, Waltham, MA)] was diluted 1:500 in 1% BSA and cells were incubated overnight at 4°C. After washing three times with PBS on a shaker, the slide was incubated with Biotinylated Secondary antibody (Vectastain Kit, Peroxidase Rabbit IgG, PK-4001) for one hour at R.T. Cells were washed three times with PBS, ABC reagent (prepared 30 minutes in advance) was added and samples were incubated for 30 minutes at R.T. After three washes with PBS, samples were incubated for 1.5 minutes with Vector Laboratories DAB Peroxidase (HRP) Substrate Kit (NC9276270. Vector laboratories, Burlingame, CA). Cells were counterstain with hematoxylin (10 seconds, R.T.) (GHS316. Sigma-Aldrich) and mounted with SecureMount (Fisher HealthCare™ PROTOCOL™ Mounting Media. #022-208. Fisher Scientific. Thermofisher Scientific).

### Cell membrane labeling with testosterone-BSA-FITC

WM46 were seeded over coverslips in p12-well culture plate, after 24 hours cells TU2% media was removed, and cells were grown in Serum-free TU media for 24 hours. Cells were treated with APA (8mM) or DMSO as vehicle control. After one hour, cells were treated with 0.25μM testosterone 3-(O-carboxymethyl) oxime:BSA-fluorescein isothiocyanate (T-BSA-FITC) conjugate (T5771, Sigma-Aldrich, Munich, Germany) diluted in Tris-Buffer (pH 7.2) for 20 min at room temperature. Negative control cells were incubated with 0.25 μM BSA-FITC (A9771, Sigma-Aldrich, Munich, Germany) diluted in Tris-Buffer (pH 7.2) for 20 min at room temperature. The medium was then aspirated, and cells were fixed for 7 minutes with 4%PFA. Cells were washed three times with PBS (5 minutes each) and mounted with ProLong Gold antifade reagent with DAPI (#P36935; Thermo Fisher Scientific). Images were captured using a Leica DM IL microscope and registered using LAS software.

### Quantitative RT-PCR

RNA was extracted using RNeasy kit (Qiagen. #74104) following the manufacturer’s instructions. cDNA was obtained using High Capacity cDNA Reverse Transcription Kit (Applied Biosystems #4368814). For quantitative real-time PCR, PowerUP™ SYBR™ Green Master Mix (Applied Biosystems #A25741) was used. ViiA 7 Real-Time PCR System was used to perform the reaction (Applied Biosystems). Values were corrected by β-actin expression. The 2^−ΔΔCt^ method was applied to calculate the relative gene expression. Primers used for the amplification are included in Table S2.

### Mice, subcutaneous tumors and pharmacologic treatments

All mice were purchased from Taconic Biosciences, Inc. (Rensselaer, NY, USA). 8-to 10-week old male and female C57BL/6NTac or IcrTac:ICR-Prkdcscid mice were allowed to reach sexually mature ages or to acclimatize for one week prior to being used in experiments. These studies were performed without inclusion/exclusion criteria or blinding but included randomization. Based on a two-fold anticipated effect, we performed experiments with at least 5 biological replicates. All procedures were performed in accordance with International Animal Care and Use Committee (IACUC)-approved protocols at the University of Pennsylvania. Subcutaneous tumors were initiated by injecting tumor cells in 50% Matrigel (Corning, Bedford, MA, USA) into the subcutaneous space on the right flanks of mice. 10 × 10^6^ human WM46 or 10 × 10^5^ murine YUMM1.7 cells were used for each tumor. Oral administration of vehicle, bicalutamide (30mg/kg/day) or apalutamide (20mg/kg/day) was performed daily. 100μl of drug was administrated by oral gavage [10%DMSO:90%Vehicle (15% ethanol:85% sesame oil)]. Each experiment include n=5 replicates, which provide 100% power to detect at least a 50% difference between groups with 95% confidence. The size of each animal cohort was determined by estimating biologically relevant effect sizes between control and treated groups and then using the minimum number of animals that could reveal statistical significance using the indicated tests of significance. When in vivo experiments were performed with female, mice located in different cages, animals were randomized prior to cell inoculation or drug treatment. All animals housed within the same cage were placed within the same treatment group. Weight and health status of the mice as well as tumor growth were monitored daily. As subcutaneous tumors grew in mice, perpendicular tumor diameters were measured using calipers. Volume was calculated using the formula L × W^2^ × 0.52, where L is the longest dimension and W is the perpendicular dimension. Animals were euthanized when tumors exceeded a protocol-specified size of 15 mm in the longest dimension. Secondary endpoints include severe ulceration, death, and any other condition that falls within the IACUC guidelines for Rodent Tumor and Cancer Models at the University of Pennsylvania.

### Statistical analysis

For experiments comparing two groups (treated vs control), significance was calculated using the Mann-Whitney test. For experiments with more than two groups, one-way ANOVA with Tukey’s honest significance difference test was performed. For the tumor growth studies *in vivo*, non-linear regression analysis was performed to get the Exponential Growth Equation and doubling times for tumor growth. To analyze the slopes of the curves and to compare them between groups, linear regression analysis was performed. All the statistical analyses in this work were performed using Graphpad Prism Software. Error bars represent standard error of the mean (SEM). **** p value≤0.0001; *** p value≤0.001; ** p value≤0.01; * p value≤0.05; n.s>0.05.

## Results

### Melanoma tumors grow more quickly in male vs. female mice

To test whether preclinical melanoma models recapitulate the male vs. female survival disparity observed in humans, we first determined growth of human BRAF-driven melanoma (WM46, female origin, BRaf^V600E^; CDK4^R24C^). Tumors grew faster in immunodeficient SCID male mice compared with matched female mice (Fig. 1A and Fig. S1A), indicating that the differences do not depend on T and/or B cell immune responses. To test whether this phenotype extends to a genetically-engineered preclinical mouse model, we used murine YUMM1.7 cells (male origin, BRaf^V600E/wt^;Pten^-/-^CdkN2a^-/-^)^16^ in syngeneic immunocompetent C57BL/6 mice. These tumors also consistently grew faster in male mice compared with matched females (Fig. S1B).

**Fig. 1:**
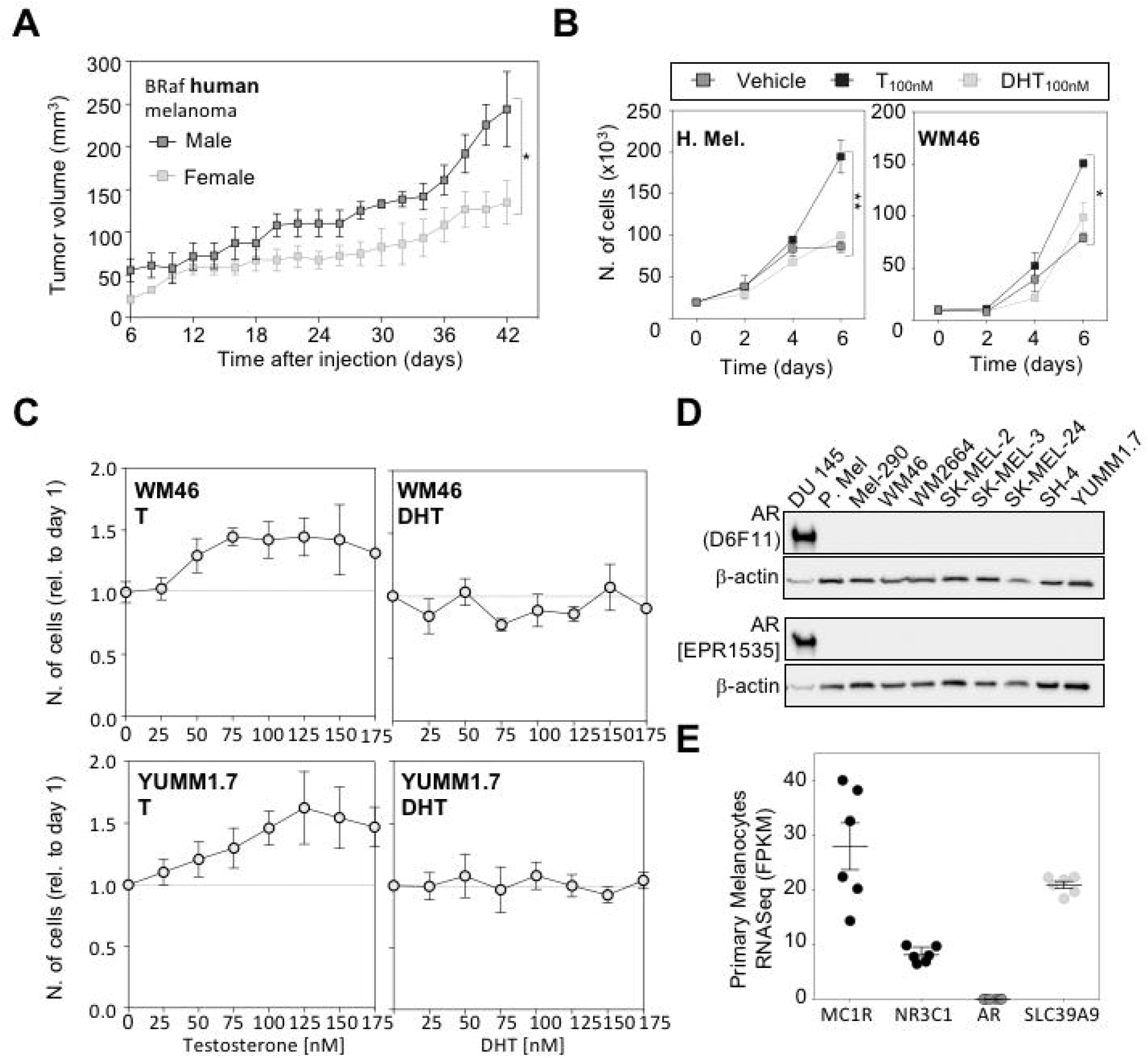
Biologic sex and testosterone promote proliferation of melanoma models lacking detectable AR. **A**. Tumor growth of human WM46 melanoma in male and female immunodeficient SCID mice. Tumor doubling times are 16.99 and 19.95 days for males and females respectively (Non-linear regression analysis/Exponential fit. See also Sup. Fig. 1 for expanded statistical analysis). **B**. Cell proliferation (cell number) determined after 6 days of treatment with vehicle (DMSO), 100nM testosterone (T) or 100nM dihydrotestosterone (DHT). Human primary melanocytes (H.Mel.) and human melanoma WM46 cells are shown. Graphs represent the average of three independent experiments. **C**. Cell proliferation of human WM46 and murine YUMM1.7 melanoma cells exposed to increasing concentrations of testosterone (T) (left panels) or dihydrotestosterone (DHT) (right panel). **D**. AR protein expression determined by Western blot with two different antibodies. Upper line = Androgen Receptor [(D6F11) XP^®^ Rabbit mAb #5153. Lower line = Recombinant Anti-Androgen Receptor antibody [EPR1535(2)] (ab133273). The prostate cancer cell line DU 145 was used as a postivie control.). β-actin was used as loading control. A replicate with increased exposure time is shown in Fig. S1G. **E**. RNA-seq data from 6 different primary melanocyte cell lines reported as median FPKM (number Fragments Per Kilobase of exon per Million reads). Melanocytic Receptor 1 (MC1R) and Glucocorticoid Receptor (NR3C1) were used as positive controls for membrane and nuclear receptors respectively. ** p value≤0.01; * p value≤0.05.

### Testosterone, but not dihydrotestosterone, promotes proliferation in melanoma cells independent of the classical androgen receptor

Physiologic concentrations of testosterone promoted proliferation of both normal early passage primary human melanocytes and melanoma cells (Fig. 1B, C and Fig. S1C). The proliferative response to testosterone correlated with dose and was saturable, suggesting a specific receptor-mediated activity. In contrast, dihydrotestosterone (DHT) had no effect on proliferation of these cell types across a wide concentration range (Fig. 1C). Dihydrotestosterone is a more potent AR agonist than testosterone^17,18^. The fact that the melanocytes and melanoma cells responded to testosterone, but not to DHT (Fig. 1B, 1C, Fig. S1C, S1D and S1E), suggests the possibility that testosterone effects in these cells are mediated by the nonclassical androgen receptor ZIP9, as ZIP9 has a much higher affinity for testosterone than for dihydroteststerone^15^. Further suggesting the idea that a nonclassical androgen receptor mediates these observed testosterone effects, we were unable to detect AR protein (via western blotting using 2 different AR antibodies, or via immunofluorescence) in primary melanocytes, or in any of the 8 melanoma cell lines we have tested (Table S1). We did, however, readily detect AR protein in 2 prostate cancer lines used as positive controls (Fig. 1D and Fig. S1F and S1G). Consistent with this lack of detectable AR protein, AR transcript was not detected via RNASeq in any of 6 different primary human melanocyte cultures. In contrast, *SLC39A9*/ZIP9 transcript, as well as transcript for the classical nuclear glucocorticoid receptor *NR3C1*, and for the melanocyte marker *MC1R*, were each readily detected in all the cell lines (Fig. 1E).

To test whether testosterone effects in melanoma cells are mediated by a surface receptor, we next treated WM46 and SK-MEL-24 human melanoma cells, (female and male derived, respectively) with a membrane-impermeable testosterone analogue [(Testosterone 3-(O-carboxymethyl)oxime: BSA] that is incapable of activating AR (AR is necessarily active in the nucleus). This compound and free testosterone each similarly promoted melanoma proliferation (Fig. S2A), supporting the idea that a non-classical androgen receptor located in the cellular membrane was responsible for the observed phenotype.

ZIP9 is broadly expressed in most cancers (Fig. S2B), normal human melanocytes, and all melanoma lines that we tested (Fig. 2A). ZIP9 transports zinc (Zn^++^) across cell and organelle membranes into the cytoplasm and is the only steroid receptor known to directly regulate zinc homeostasis^19^. To test whether testosterone activates this ZIP9 activity in melanoma, we used the fluorescent Zn^++^-specific probe FluoZin-3. Testosterone induced a rapid increase in cytosolic free Zn^++^ ([Zn^++^]_i_) in human melanoma cells (Fig. 2B), that was followed by a sustained elevation for at least 2 days in the presence of testosterone (Fig. 2C). We determined that Zn^++^ influx was necessary for the testosterone-dependent increase in melanoma cell proliferation, as treatment with the Zn^++^ chelator N,N,N′,N′-tetrakis (2-pyridinylmethyl)-1,2-ethanediamine (TPEN), blocked testosterone induced proliferation at TPEN concentrations that had no significant effect on proliferation when used alone (Fig. 2D and Fig. S2C).

**Fig 2:**
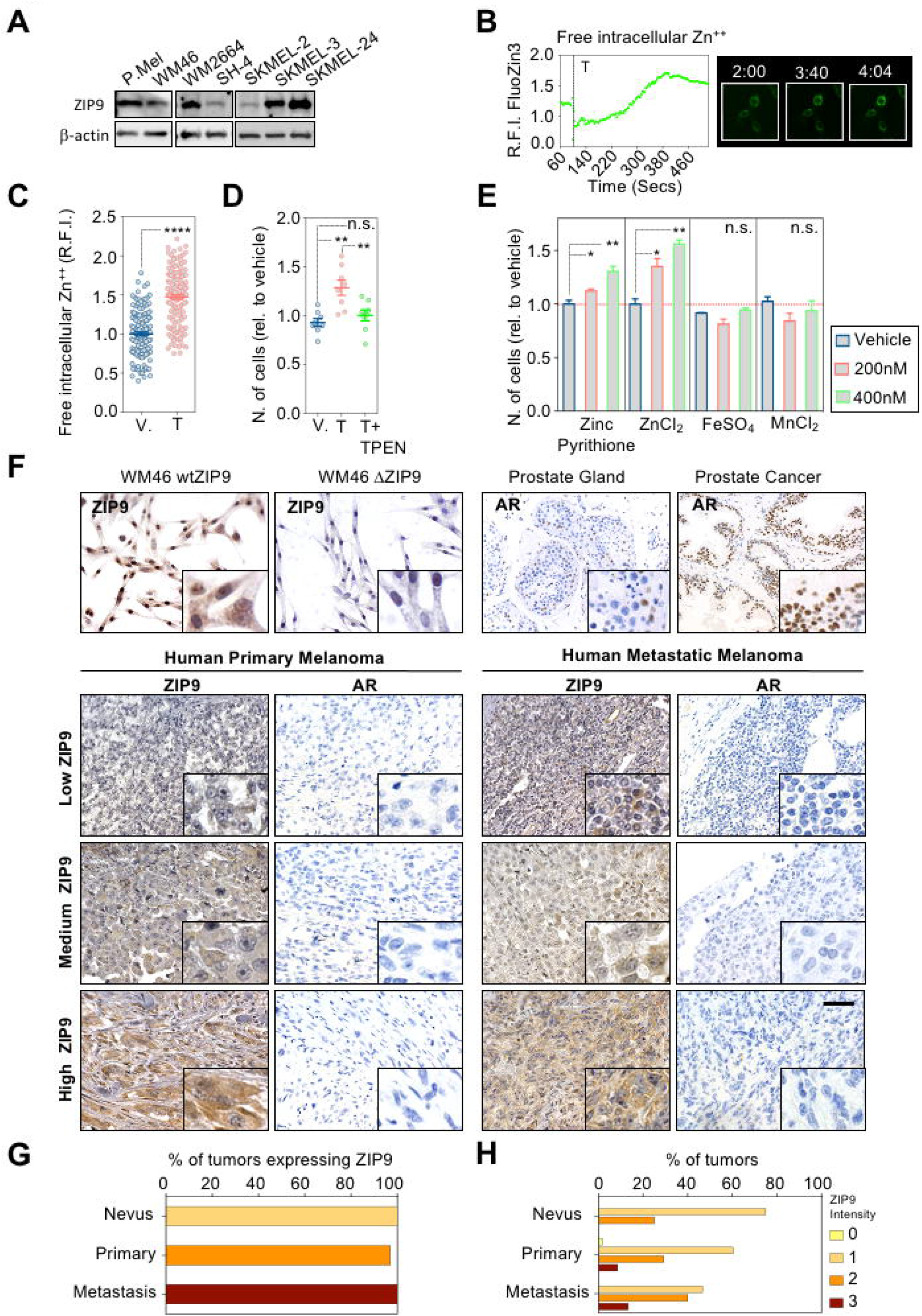
ZIP9 is active in human melanoma cells, and is broadly expressed in human melanocytic tumors. **A**. ZIP9 protein expression determined by western blot in primary melanocytes and a battery of human melanoma cell lines. **B**. Time-lapse *in vivo* analysis of Zn^++^influx in WM46 cells upon testosterone addition (100nM). FluoZin-3 was used as Zn^++^ reporter. **C**. Intracellular levels of Zn^++^ after long-term testosterone treatment (96 hours; 100nM testosterone). Zinc levels were measured as fluorescence intensity per cell. FluoZin-3 was used as Zn^++^ reporter. Representative images are shown on the right at indicated time-points. **D**. WM46 relative proliferation (cell number) after 6 days in the presence of 100 nM testosterone (T) +/-200 nM zinc chelator (TPEN). **E**. WM46 proliferation in the presence exogenous divalent cations Zn^++^ Fe^++^ and Mn^++^. Cells were grown for 6 days and treated as indicated in the legend. Error bars represent standard error of the mean (SEM). **F**. Validation of ZIP9 [(SLC39A9 Antibody (PA5-52485)] and androgen receptor [(Leica AR-318-L-CE, clone AR27 (clone AR27, 1:25)] antibodies for immunohistochemistry. ZIP9 staining performed in wild-type and ZIP9 knock-out cells. Non permeabilization of the cells ensured ZIP9 membrane localization. Prostate gland tissue and human prostate cancer samples were used as positive controls for AR. Representative images of human melanoma samples stained for ZIP9 and AR. Tumors expressing low, medium and high levels of ZIP9 are shown. Replicates from the same samples stained for AR are shown. 20X magnification (1.6X zoom). Scale bar=60μM. **G**. Graphic representation of the % of tumors that express ZIP9. Data from nevus, primary melanomas and metastatic melanoma are displayed. **H**. Graphic representation of the percentage of nevi, primary lesions and metastatic tumors classified according to ZIP9 intensity (Score 1=1-25%, 2=26-50%, 3=51-75%, 4=76-100%). **** p value≤0.0001; ** p value≤0.01; * p value≤0.05; n.s>0.05.

Although TPEN has high affinity for Zn^++^, it also has some ability to chelate copper^20^. To test whether copper flux might contribute to testosterone effects on melanoma proliferation, we treated WM46 cells with CuSO_4_, which had no significant effect on cell proliferation (Fig. S2D). Moreover, the highly specific copper chelator bathocuproinedisulfonic acid (BCS), did not affect the proliferative response to testosterone, nor did BCS affect testosterone induced Zn^++^ influx (Fig. S2E and Fig. S2F).

After determining that Zn^++^ influx was necessary for testosterone-induced proliferation, we next tested whether increased Zn^++^ was sufficient to increase melanoma proliferation. Exogenous zinc pyrithione (also a Zn^++^ ionophore), and ZnCl_2_ each increased melanoma proliferation (Fig. 2E). Other biologically-relevant divalent cations, Fe^++^ and Mn^++^, known to be transported by other ZIP family members^21^, had no effect on proliferation (Fig. 2E). Together, these results demonstrate that testosterone, but not DHT, promotes zinc dependent proliferation in melanoma cells that express ZIP9 and lack detectable AR.

We next used immunocytochemistry to test whether AR and/or ZIP9 protein are expressed in human melanocytic lesions (14 benign nevi, 63 primary melanomas, and 21 metastatic melanomas from both males and females). AR was determined in tissue sections using a highly validated CLIA (Clinical Laboratory Improvement Amendments) certified method in the clinical pathology lab at the Hospital of the University of Pennsylvania. This procedure uses a validated antibody different from the two we used for western blotting and immunofluorescence. Each tissue was scored in a blinded fashion by a board-certified pathologist. While AR was readily detectable in prostate tissue used as positive control, AR was not detected in any of the nevi, nor in the melanomas (Fig. 2F). In parallel, we analyzed the same samples for ZIP9. CLIA grade ZIP9 IHC is not available. However, we validated our ZIP9 antibody using parental WM46 ZIP9 positive (wtZIP9) and isogenic ZIP9 negative (ΔZIP9) cell lines. These cells were grown on chamber slides and processed for IHC in parallel with the human samples. The antibody used labeled only the wtZIP9 cells. ZIP9 protein was observed in 100% of the nevi, 97% of primary melanomas and 100% of the metastatic samples (Fig. 2G). Further, the ZIP9 relative staining intensity positively correlated with tumor stage (Fig. 2H). ZIP9 intensity was skewed toward higher intensity scores (Score 2 and 3) in metastatic tumors (Score 0: 0%; Score 1: 46.6%; Score 2: 40%; Score 3: 13.3% of tumors) vs. primary tumors (Score 0: 1.72%; Score 1: 60.34%; Score 2: 29.31%; Score 3: 8.62% of tumors. None of the nevi showed the highest level (Score 3) observed in the melanomas. No significant differences in ZIP9 staining were observed between males and females (Fig. S2G). Similarly, no correlation was observed between ZIP9 expression and the age of the patient (Fig. S2H).

ZIP9 is broadly expressed and its pro-melanoma activity is likely limited more by the availability of testosterone, than by ZIP9 itself. However, we did question whether ZIP9 expression correlates with clinical melanoma outcomes in people. We used *OSskm*^22^ to analyze pooled data from 1085 tumors (563 death events) from multiple studies (GSE17275, GSE19234, GSE22155_GPL6102, GSE22155_GPL6947, GSE50509, GSE46517, GSE53118, GSE65904, GSE98394 and TCGA. Although not significant (p-value=0.11), there is a trend associating higher ZIP9 expression with shorter survival (HR=1.2039) (Fig S3A and Table S2), consistent with the idea that ZIP9 promotes melanoma progression. Importantly, when patients are stratified by sex (TCGA), the trend between high ZIP9 expression and poor survival is observed in males (H.R.=1.27, C.I. 95% 0.64-2.52), but not in females (H.R. 1.01, C.I. 95% 0.41-2.46) (Fig. S3B). No correlation was found between ZIP9 expression and several other prognostic melanoma markers including age, tumor ulceration, Clark level, mitotic rate, and aneuploidy Fig. S3C).

### Testosterone signaling through ZIP9 promotes melanoma growth

To determine whether ZIP9 mediates testosterone effects, we used CRISPR-Cas9 and gRNA targeting Slc39A9 exon 1 to ablate ZIP9 in human (WM46) melanoma and in murine (YUMM1.7) melanoma. Isogenic clonal populations of parental ZIP9-expressing (wtZIP9) and ZIP9-ablated (ΔZIP9) cells were established, and the mutations disrupting the reading frame were mapped by sequencing (Fig. S4A, S4B and S4C). ZIP9 protein was not detectable in ΔZIP9 cells (Fig. 2F, Fig. 3A, Fig. 3B and Fig. S4D). ΔZIP9 cells did not respond to testosterone, whereas isogenic wtZIP9 control clones responded similarly to the parental wild-type cells (Fig. 3A, 3B and Fig. S4E).

**Fig. 3:**
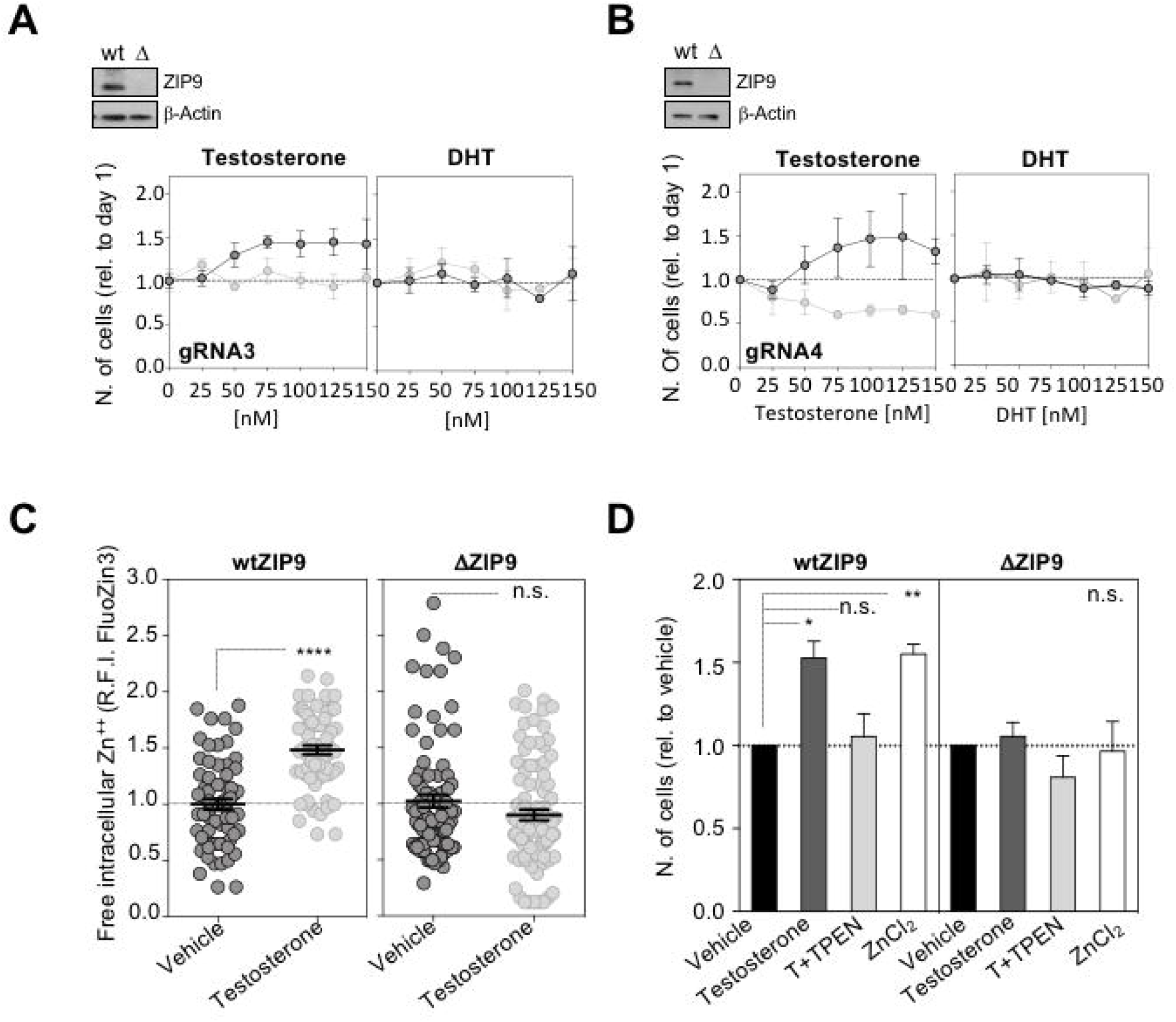
ZIP9 mediates testosterone effects in melanoma. **A**. Proliferation (cell number) of isogenic clonal populations of WM46 wtZIP9 and ΔZIP9 (gRNA #3) cells exposed to increasing concentrations of testosterone (T) and dihydrotestosterone (DHT). **B**. Proliferation (cell number) of isogenic clonal populations of WM46 wtZIP9 and ΔZIP9 (gRNA #4) cells exposed to increasing concentrations of testosterone (T) and dihydrotestosterone (DHT). The graph represents the average of three independent experiments. **C**. Intracellular levels of zinc in human melanoma ΔZIP9 cells measured as relative fluorescence intensity of FluoZin-3. Graphs represent the average of three independent experiments. **D**. Relative cell proliferation in the presence of 100nM testosterone (T), zinc chelator (200nM TPEN) and ZnCl_2_ (400nM). Cells were cultured for 6 days. The graph represents the average of three independent experiments. **** p value≤0.0001; ** p value≤0.01; * p value≤0.05; n.s>0.05.

The testosterone insensitive phenotype associated with CRISPR-Cas9 engineered ZIP9 loss was rescued by expression of a human ZIP9 transgene. Lentiviral mediated constitutive ZIP9 expression in the ΔZIP9 WM46 cells restored both ZIP9 protein and the proliferative response to testosterone, verifying the on-target effects of the CRISPR-Cas9 mediated ZIP9 ablation (Fig. S4F, S4G).

Consistent with the conclusion that ZIP9 is the major mediator of testosterone effects in these melanoma cells, exposure to testosterone failed to increase [Zn^++^]_i_ in ΔZIP9 cells (Fig. 3C and 3D). Intracellular free zinc levels following exogenous zinc exposure are similar between wtZIP9 and ΔZIP9 cells (Fig. S4H), indicating that while other zinc transporters are expressed in human melanoma cells, ZIP9 induces a rapid, ZIP9-dependent, increase in [Zn^++^]_i_ that is required for the increased proliferation in melanoma cells (Fig. 3D and Fig. S4H).

### Testosterone-induced melanoma proliferation through ZIP9 requires downstream MAPK and YAP1 activation

Although the studies detailed above show that testosterone promotes melanoma proliferation via ZIP9 dependent Zn^++^ influx, zinc is involved in myriad cellular processes, making it difficult to predict *a priori* which downstream signaling pathways are required for the testosterone activity. To start identifying these, we used a 447-element Reverse Phase Protein Array (RPPA) analysis of human WM46 melanoma cells treated with testosterone for 0, 30, 60 and 480 min. While the relative expression of most represented proteins was unaffected by testosterone, some were significantly under or overexpressed, including several with known tumor-promoting or tumor-suppressive functions (Fig. 4A and Table S3). Downregulated proteins included 14-3-3ε, a tumor suppressor and negative YAP1 regulator previously implicated in liver, lung, and gastric cancers^23^, and CDKN2A (p16), a cyclin-dependent kinase (CDK) inhibitor, and one of the most studied tumor suppressors^24^. Upregulated proteins included key elements of tumor-promoting pathways, most notably phosphorylated ERK (T202; Y204) and YAP1 (Fig. 4A). Importantly, ΔZIP9 cells displayed decreased basal levels of ERK phosphorylation compared with isogenic wild-type clones (Fig. S5A). MAPK activation was necessary for the testosterone-dependent proliferative response, as the specific ATP-competitive ERK1/2 inhibitor, ulixertinib^25^ (RVD-523; Ei) blocked the testosterone-dependent proliferative response, at ulixertinib concentrations that had no effect on the basal proliferation rate when used alone (Fig. 4B). Testosterone-dependent ZIP9 activation appeared to render WM46 cells more sensitive to ulixertinib, as cell viability was compromised when they were treated with both testosterone and RVD-523. This vulnerability to combination treatment seems to be specific and ZIP9 dependent, as ΔZIP9 cells treated with both testosterone and ulixertinib proliferated at rates comparable to controls (Fig. 4B).

**Fig. 4:**
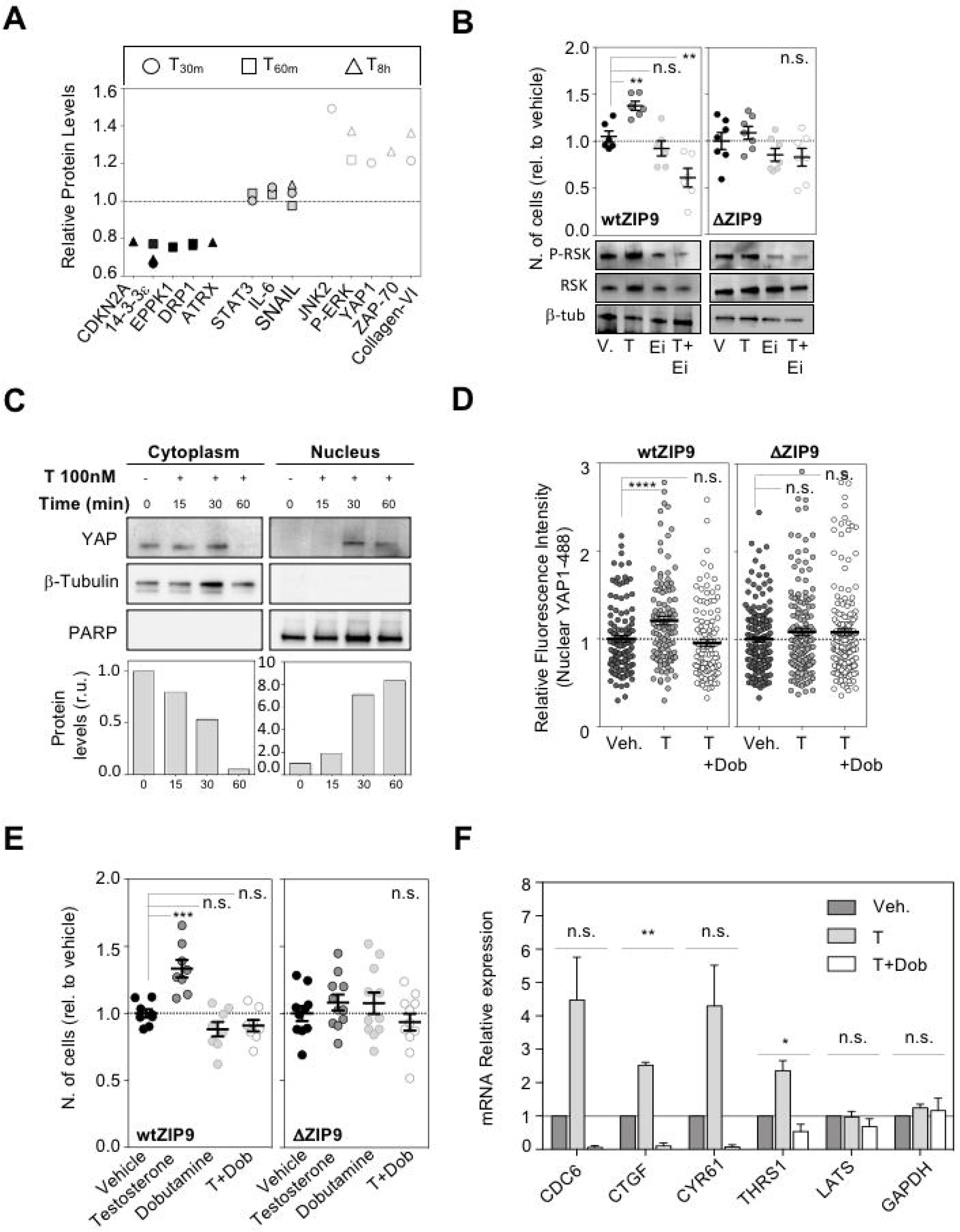
Testosterone driven increase in melanoma proliferation requires ZIP9 and activation of MAPK and YAP1. **A**. RPPA analysis displaying changes in protein expression in WM46 human melanoma cells following exposure to 100 nM testosterone (T) for increasing amounts of time. Down-regulated proteins are shown in black; white color corresponds to up-regulated proteins and control proteins showing no fold-change when compared to vehicle-treated cells are shown in dark grey. **B**. Relative proliferation (cell number) after exposure to pharmacologic ERK1/2 inhibition via 50nM RVD-523 (Ei) alone or in combination with testosterone. wtZIP9 and ΔZIP9 WM46 cells are shown. The western blot shows levels of phosphorylation of the ERK target RSK in wtZIP9 and ΔZIP9 WM46 cells (Ei represents RVD-523). **C**. Western blot for YAP1 in fractionated WM46 lysates. Cells were treated with 100 nM testosterone for the indicated times. β-Actin is used as cytoplasmic fraction positive control. PARP is used as nuclear fraction positive control. Quantification of protein levels normalized against T_0_ is shown in the histograms (lower panel) **D**. Quantification of YAP-1 nuclear immunodetection after 30 minutes of exposure to 100 nM testosterone (T) and/or 8μM dobutamine (Dob) in wtZIP9 and βZIP9 cells. **E**. Proliferation of wtZIP9 and βZIP9 WM46 cells after treatment with 100nM testosterone (T) and/or the YAP inhibitor dobutamine (Dob). **F**. Relative mRNA expression of YAP1 target genes after 30 minutes in the presence of 100 nM testosterone and/or 8μM dobutamine. Error bars represent standard error of the mean (SEM). *** p value≤0.001; ** p value≤0.01; * p value≤0.05; n.s>0.05.

We determined that YAP1 activation is also required for the augmented proliferative response driven by testosterone. YAP1 is a transcriptional coactivator whose activity is largely regulated by its localization^26^. Testosterone induced rapid YAP1 translocation from cytoplasm to nucleus in human melanoma cells in a ZIP9-dependent manner (Fig. 4C, 4D and Fig. S5B). YAP1 subcellular localization is controlled largely by LATS, which phosphorylates YAP1 at Serine 127 and thereby retains YAP1 in the cytoplasm^27^. The YAP1 inhibitor dobutamine also promotes phosphorylation of YAP at Ser127^28^. Consistent with this, the testosterone induced YAP1 nuclear localization and increase in cell proliferation were both blocked by dobutamine, while this compound had no significant effect on its own (Fig. 4D, and Fig. 4E). When ZIP9 expression was rescued via lentiviral transduction of a ZIP9 transgene to ΔZIP9 WM46 cells, testosterone dependent YAP1 translocation into the nucleus was also restored (Fig. S5C).

Testosterone promotion of YAP1 activity was evidenced by increased expression of several well-known YAP1 target genes including CDC6, CTGF, CYR61 and THBS1. As expected, these testosterone-induced expression changes were also blocked by dobutamine (Fig. 4F).

### Pharmacologic ZIP9 blockade inhibits testosterone-driven melanoma proliferation and melanoma tumor growth *in vivo*

We next tested whether FDA-approved drugs could be repurposed to effectively target ZIP9. Although specific ZIP9 inhibitors are not yet developed, molecular modeling and competition assays with fluorescent-tagged testosterone suggest that the androgen receptor inhibitor bicalutamide (BIC) competes with testosterone for ZIP9 binding to the same extracellular pocket and thereby acts as a competitive ZIP9 antagonist^29^. Bicalutamide was developed and approved for prostate cancer and has now been largely replaced by enzalutamide (ENZ) and apalutamide (APA), which are structurally related analogs with higher affinity for the androgen receptor and greater clinical efficacy against advanced prostate cancer^30,31^.

While ZIP9 was not known to be an androgen receptor at the time these drugs were developed, we show here that they are nonetheless effective inhibitors of testosterone effects in melanoma cells that express ZIP9, but that lack detectable AR. Each compound completely blocked testosterone-induced proliferation and MAPK activation in several melanoma cell lines (Fig. 5A, 5B). Importantly, these agents alone (without testosterone) had no effect on cell proliferation (Fig. S6A). The testosterone dependent increase of intracellular zinc was also efficiently blocked by bicalutamide (Fig. S6B). Moreover, ΔZIP9 cells transduced to re-express ZIP9, responded to testosterone, and this effect was again blocked by bicalutamide (Fig. S6C).

**Fig. 5:**
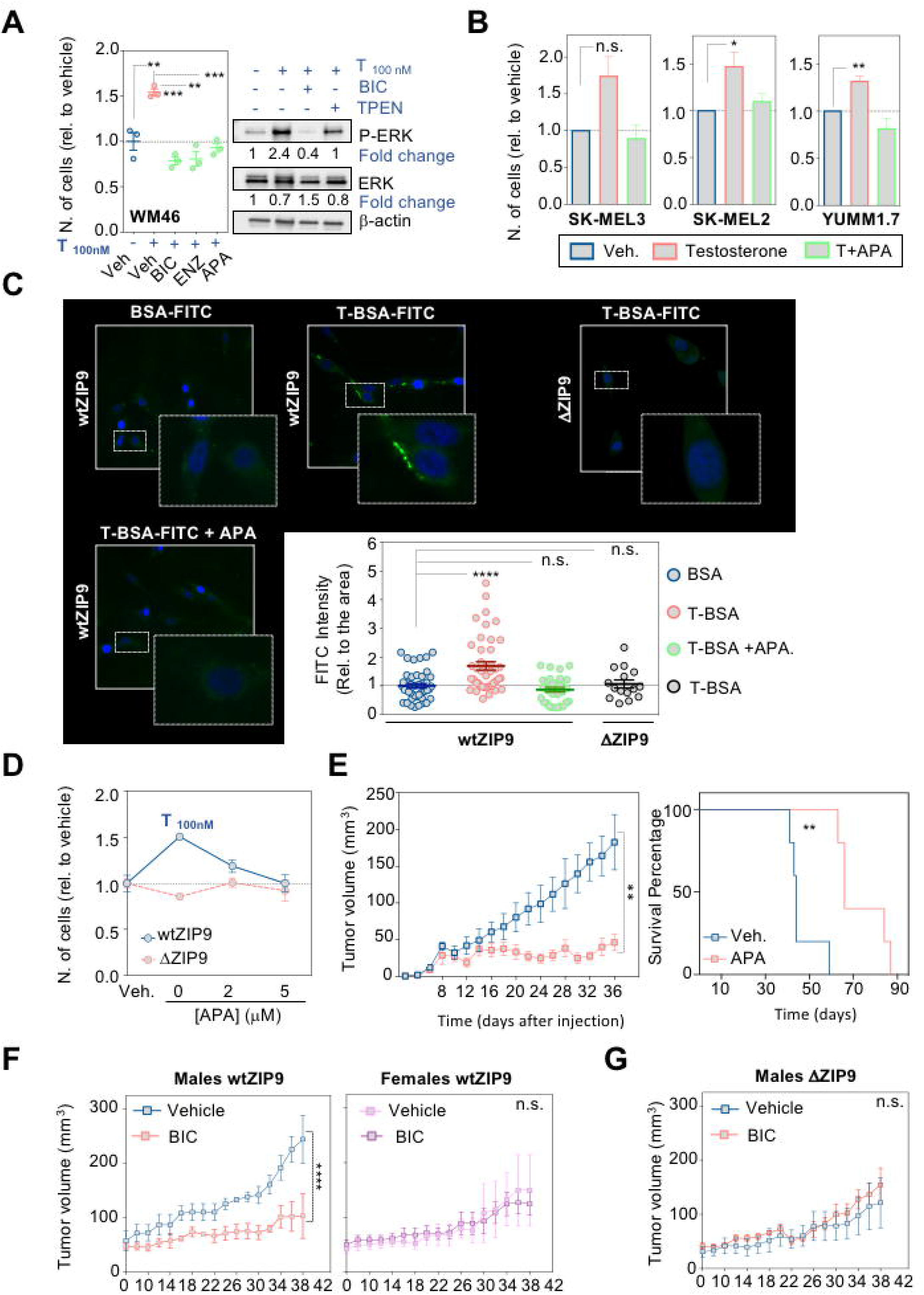
Pharmacologic ZIP9 blockade inhibits melanoma *in vivo*. **A**. Proliferation of human melanoma cells (WM46) in the presence of 100 nM testosterone (T) +/-2 μM AR inhibitors (BIC:Bicalutamide; ENZ:Enzalutamide; APA:Apalutamide). Western blot showing ERK and p-ERK proteins in WM46 cells treated with 100nM testosterone +/-2μM BIC +/-200 nM zinc chelator (TPEN). **B**. Cell proliferation in human and murine derived melanoma cells [SK-MEL-3 (male), SK-MEL-2 (female) and YUMM1.7(male)] treated with 100 nM testosterone (T) in combination with apalutamide (2 μM) (APA). **C**. Cell membrane labeling with cell impermeable testosterone-BSA conjugated with FITC (0.25 μM). BSA-FITC (0.25 μM) was used as a negative control for unspecific binding. Quantification of membrane labeling with T-BSA or the control BSA. The graph represents the fluorescence intensity relative to the total area of each cell. **D**. Proliferation of WM46 wtZIP9 and Δ ZIP9 treated with 100 nM testosterone (T) in combination with apalutamide (2 μM) (APA). **E**. Tumor growth and survival analysis in SCID male mice bearing WM46 derived subcutaneous tumors (APA treatment: 20 mg/kg/day via oral gavage). ** p-value<0.005 by ANOVA. Doubling time (non-linear regression analysis): Vehicle=11.11 days; APA=25.22 days. **F**. Tumor growth in mice bearing WM46 derived subcutaneous tumors. Daily treatment with bicalutamide (30 mg/kg/day, via oral gavage) or vehicle are shown for both male and female mice. Linear regression analysis of slopes demonstrates significant differences between vehicle treated and Bic treated males (p-value<0.0001) (See Fig. S6B and S6C). **G**. Tumor growth in SCID male mice bearing ΔZIP9 WM46 melanoma. Mice were treated daily with bicalutamide (30 mg/kg/day) or vehicle via oral gavage. See also Sup. Fig. 6 for expanded statistical analysis). **** p value≤0.0001; *** p value≤0.001; ** p value≤0.01; * p value≤0.05; n.s>0.05.

To further confirm that these pharmacologic agents work through ZIP9, we next used cyproterone acetate (CPA), an anti-androgen that blocks the testosterone interaction with AR, but that does not bind to ZIP9^32^. In our melanoma cells, up to a 20-fold molar excess of CPA did not significantly inhibit testosterone effects on proliferation, consistent with the conclusion that AR is not the major mediator of testosterone effects in these melanoma models (Fig. S6D). Importantly, response to testosterone and/or APA was independent on BRAF status, as SK-MEL-2 cells (B-RAF^wt^) responded well to testosterone, non-permeable testosterone, and to ZIP9 inhibition by apalutamide (Fig S6E).

BRAF inhibitors are useful in melanoma patients with BRAF driven tumors ^33,34^. To test whether combined targeted inhibition of BRAF and ZIP9 might be effective against BRAF mutant melanoma we determined proliferation of human WM46 (BRAF^V600E^) and murine YUMM1.7 (BRAF^V600E^) in the presence of the BRAF inhibitor PLX-4032 alone, and in combination with apalutamide. Although addition of apalutamide to PLX-4032 was not synergistic in WM46 cells (possibly because PLX-4032 alone was quite effective at limiting proliferation), the combination of apalutamide and PLX-4032 was more effective in YUMM1.7 compared to either agent alone (Fig. S6F).

To test whether bicalutamide class AR inhibitors block the physical interaction between testosterone and ZIP9 in melanoma, we performed direct binding assays using a membrane-impermeable testosterone analogue (T-BSA-FITC). This reagent labels the plasma membrane surface of wild-type ZIP9 expressing WM46 melanoma cells (Fig. 5C). This membrane bound testosterone was displaced by apalutamide, demonstrating the specificity of the interaction (Fig. 5C). Further demonstrating that the binding is ZIP9 specific, testosterone localization at the plasma membrane was markedly reduced in ΔZIP9 cells (Fig. 5C) that did not respond to testosterone, nor to APA *in vitro* (Fig. 5D).

Next, we tested whether apalutamide and related analogs inhibit melanoma *in vivo*. For this, we introduced wtZIP9 or isogenic ΔZIP9 human WM46 melanoma into SCID mice. Once-daily systemically administered apalutamide (20 mg/kg/day via oral gavage) significantly suppressed growth of wtZIP9 tumors in male mice and extended survival (doubling time for tumor growth was 11.1 days for vehicle treated males and 25.2 days for APA treated males) (Fig. 5E, Fig. S7A). Similar results were obtained when males were treated with bicalutamide (30mg/kg/day) (Fig. 5F). Importantly, bicalutamide had no effect on wtZIP9 tumors in female mice, nor on ΔZIP9 tumors (Fig. 5F, G, Fig. S7B and C). Together, these data show that ZIP9 promotes melanoma progression specifically in males.

## Discussion

High levels of circulating testosterone are associated with increased risk of early death after a cancer diagnosis in men and women, and reaches 1.52 (1.20–1.91) (20.2-51.2 nmol/l) for the highest quintile.^35^ Together with this, other recent work correlating testosterone level with the risk of 19 types of cancer ^9^ shows that higher free and total testosterone in men is associated with higher risk of melanoma [free testosterone: 1.35 (95% CI 1.14, 1.61), total testosterone: 1.28 (95% CI 1.05, 1.55)] however, no association between testosterone and melanoma was found in women [free testosterone: 0.96 (95% CI 0.77, 1.20), total testosterone: 0.95 (95% CI 0.82, 1.10)]. These epidemiological data support the idea that that testosterone promotes melanoma. Outcomes for males are likely further worsened by the fact that they lack female levels of estrogen, which is increasingly recognized to play protective roles against melanoma^6,36,37^.

Many have speculated how sex steroids contribute to sex differences in cancer pathobiology, however, definitive functional studies are lacking, and the mechanisms by which they contribute to the male-female cancer sex gap are only now emerging. The role of nonclassical estrogen, progesterone, and testosterone receptors (GPER1, PAQR7, and ZIP9, respectively) in cancer pathobiology has been largely unexplored. However, these are highly likely to be significant contributors to sex specific pathobiology in melanoma (and other cancers) as each are transcribed at much higher levels than the classical nuclear sex steroid receptors, most of which are not detectable above background (Fig. S8A).

Circulating testosterone in humans generally ranges between 8-35nM^38^. We used 100 nM in our *in vitro* experiments, a concentration used in diverse cell types in different labs ^10,39,40^. For biologically active small molecules including drugs, hormones, and ions, differences in IC_50_ and saturating concentrations are often observed between *in vitro* and *in vivo* conditions. This results from differences in compound stability and protein binding which affect half-life and the amount of biologically available free vs. bound compound. These pharmacokinetic differences typically make direct comparisons between *in vitro* and *in vivo* settings difficult. Many endogenous ligands like testosterone, are only replenished in culture when the media is changed, whereas homeostatic mechanisms keep testosterone levels stable *in vivo*. A higher starting concentration *in vitro* may therefore be needed to maintain saturating levels for the entire time period between media changes.

Testosterone promoted proliferation of melanoma cells in a saturable and dose-dependent manner. Curiously, the same cell lines did not respond to DHT, which is generally considered the more biologically active androgen. DHT displays 4-fold increased affinity than testosterone for AR, and it dissociates from AR three time slower than testosterone^18^. Lack of response to DHT in melanoma is consistent with the absence of detectable AR expression in the models used for this study, and is highly consistent with our genetic data showing that ZIP9, which has higher affinity for testosterone vs. DHT, is an important mediator of testosterone effects. Consistent with this, a previous study in metastatic prostate cancer suggested that activation of the migratory machinery depends solely in the activation of testosterone/ZIP9 pathway and not in the interaction of this androgen with the nuclear AR^41^.

We show here that ZIP9 activation, via testosterone binding, promotes an increase in cytosolic zinc in melanoma cells. The mammalian family of zinc transporters SLC39A comprises 14 members (ZIP1–14)^42^ grouped into four subfamilies that were stablished according to their amino acid sequence similarities. There may be roles for these other family members in some cancers, as ZIP1 (SLC39A1, from ZIPII subfamily) has been associated with the regulation of zinc uptake in prostate cancer cells^43^. However, the rapid increase of intracellular zinc through ZIP1 appears to be AR dependent, as PC-3 cells only respond to testosterone after they are transfected with exogenous AR. ZIP9 amino acid residues predicted to be most critical for testosterone and bicalutamide binding include Ala167, Val241, Met248, and Ala167, Leu302, Ser171, respectively^29^, however only Ala167 is present in ZIP1, suggesting that this receptor may not be similarly regulated by testosterone. Regarding other SLC39A proteins, sequence analysis places ZIP9 as unique member of ZIPI subfamily^44^, and it is the only one known to interact with testosterone.

There may be many drivers of the cancer sex gap in humans, including differences in immune surveillance^45^. However, differences in immune surveillance do not appear to be a major driver of the differences in the melanoma models used for this study, as tumors progressed faster not only in male vs. female syngeneic immunocompetent mice, but also in human melanomas grown in male vs. female SCID mice. Therefore, the testosterone effects on melanoma in these models are dependent on ZIP9, but independent of B and T cell mediated anti-tumor activity. Consistent with this, ZIP9 expressing tumors responded to bicalutamide and apalutamide in SCID mice.

As ZIP9 is widely expressed in nearly all tissues, it may be a major determinant of the sex disparity in outcomes not just for melanoma, but also for many other cancer types. Consistent with this, we observed that testosterone promotes proliferation of genetically diverse melanoma lines and that this effect is blocked by bicalutamide (Fig. S9).

While this work clearly establishes a major role for ZIP9 in melanoma, we do recognize the possibility that some melanoma cell lines, and perhaps even some human tumors, may express a low level of AR that also impacts melanoma. However, any such tumor would still likely also be affected by ZIP9. A recent report^46^ also considered a possible role for testosterone in melanoma and, consistent with our data, concluded that testosterone promotes melanoma progression. Authors attributed this effect to the classical AR. However, that report did not consider the nonclassical androgen receptor ZIP9. Critically, that study did not show that AR was necessary for melanoma response to pharmacologic AR inhibitors, nor did it test whether AR was a determinant of sex differences in melanoma.

The demonstration here that ZIP9 is pharmacologically accessible, suggests that ZIP9 may be a new eminently druggable therapeutic target (Fig. 6), and that currently approved androgen receptor inhibitors might be useful in combination with current standard of care therapeutics for a wide range of cancers, especially those that disproportionately affect males.

**Fig. 6:**
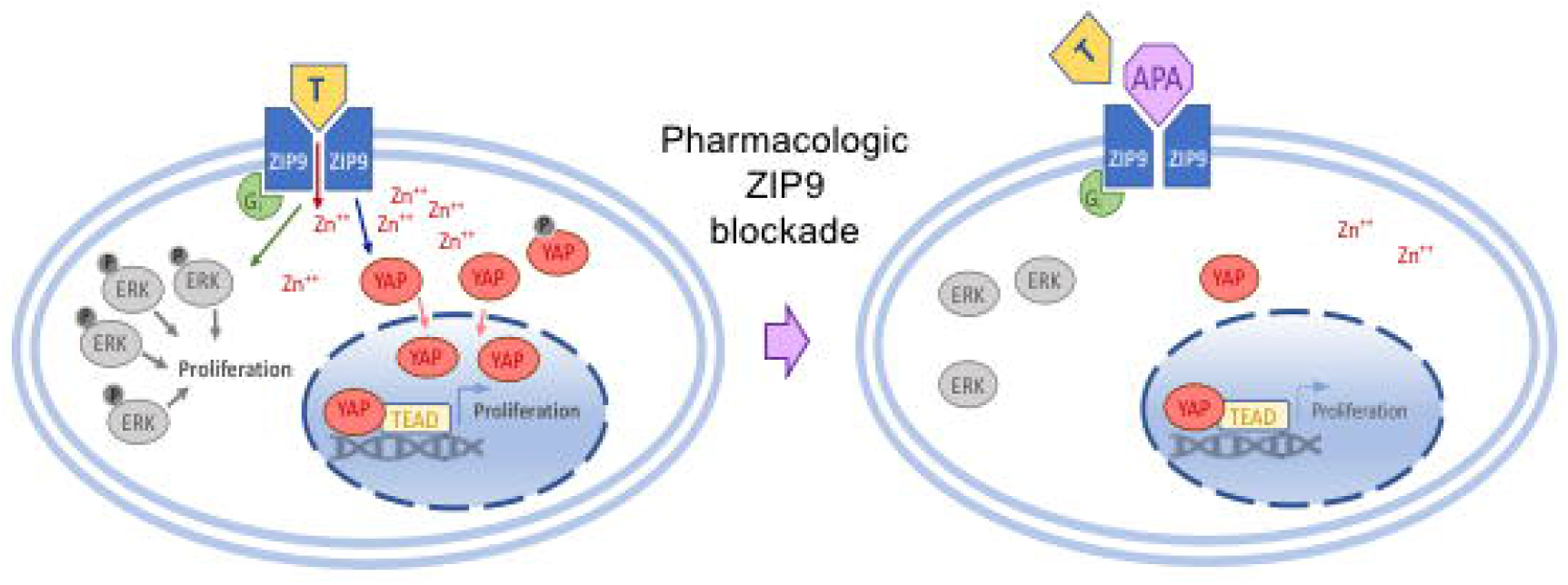
Working model. ZIP9 activation promotes ERK phosphorylation and induces YAP1 nuclear translocation. AR inhibitors [represented in the figure by apalutamide (APA)] block testosterone effects through ZIP9 inactivation.

## Acknowledgements

The authors thank the University of Pennsylvania Skin Biology and Disease Research-based center for analysis of tissue sections and University of Pennsylvania Pathology Clinical Service Center—Anatomic Pathology Division for the AR staining of the tissue microarray.

## Notes

**<u>Financial support:</u>** T.W.R. is supported by grant from the NIH/NCI (R01 CA163566, R41CA228695), and by the Stiefel award from the Dermatology Foundation. This work was also supported in part by the Penn Skin Biology and Diseases Resource-based Center (P30-AR069589), the Melanoma Research Foundation and DOD to T.W.R., and R37 GM56328 to J.K.F.

### Competing Interest Statement

C.A.P., C.A.N., and T.W.R., and are inventors on a provisional patent held by the University of Pennsylvania related to this work.

### Summary of Updates

New confirmatory data. Clarified text, discussion and data.

