## Supplementary Figure 1 for "ZIP9 is a Druggable Determinant of Sex Differences in Melanoma"

**A**

| Non linear reg. | Males | Females |
|---|---|---|
| Y0 | 41.04 | 31.11 |
| k | 0.04079 | 0.03474 |
| Tau | 24.52 | 28.79 |
| Doubling Time | 16.99 | 19.95 |
| **Std. Error** | | |
| Y0 | 4.767 | 4.464 |
| k | 0.003447 | 0.004367 |

**B**

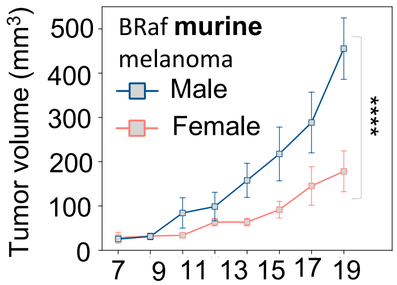


| Non linear reg. | Males | Females |
|---|---|---|
| Y0 | 8.82 | 7.748 |
| k | 0.1871 | 0.1501 |
| Tau | 5.345 | 6.663 |
| Doubling Time | 3.705 | 4.619 |
| **Std. Error** | | |
| Y0 | 4.76 | 4.195 |
| k | 0.02781 | 0.0286 |

**C**

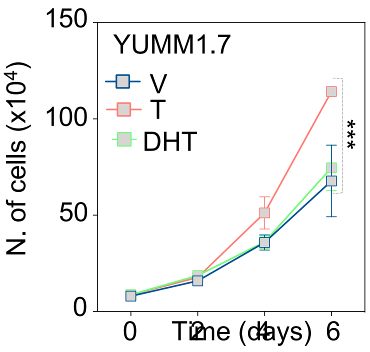


**D**

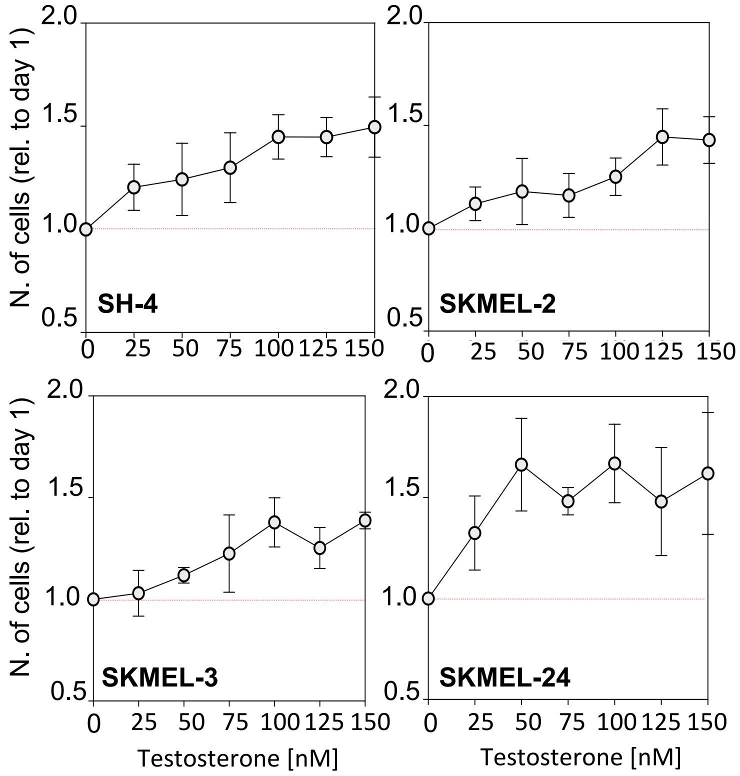


**E**

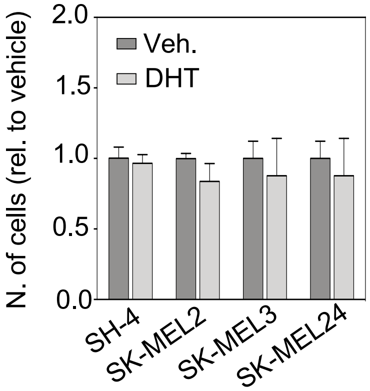


**F**

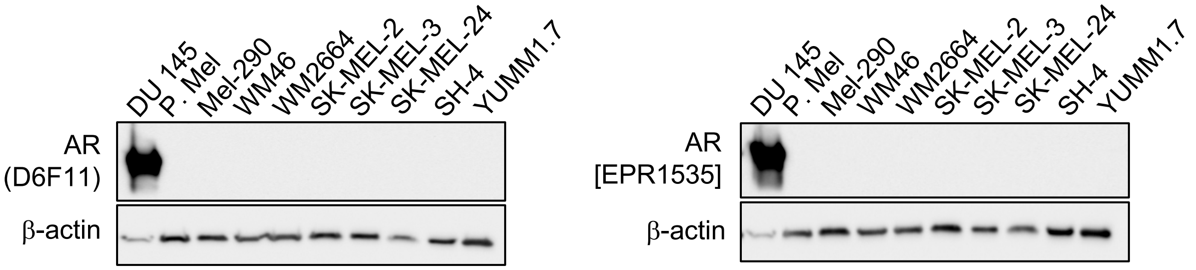


**G**

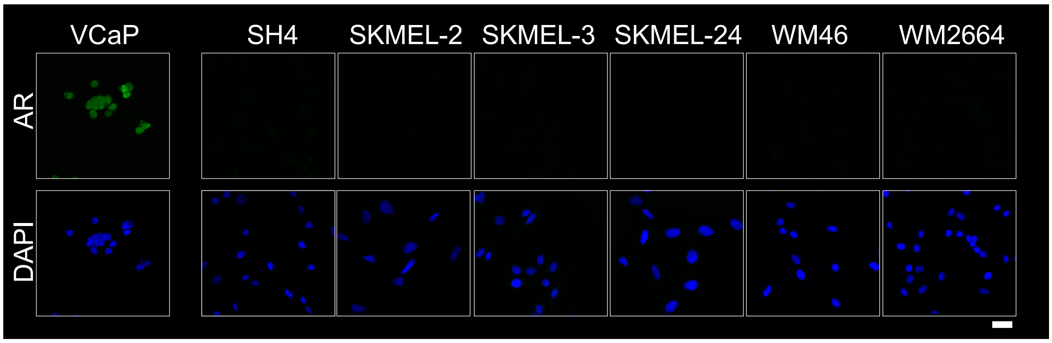
