## Supplementary Figure 4 for "ZIP9 is a Druggable Determinant of Sex Differences in Melanoma"

**A**

| ICE Analysis_Synthego | Indel % | Model Fit (R2) | Knock-out Score |
|---|---|---|---|
| WM46 wtZIP9 gRNA 3. Clone 3.3 | 0 | 1 | 0 |
| WM46 koZIP9 gRNA 3. Clone 3.2 | 92 | 0.92 | 92 |
| WM46 wtZIP9 gRNA 3. Clone 4.6 | 0 | 1 | 0 |
| WM46 koZIP9 gRNA 4. Clone 4.b | 90 | 0.92 | 90 |
| YUMM1.7 koZIP9 gRNA 3. Clone 3.3 | 90 | 0.9 | 90 |

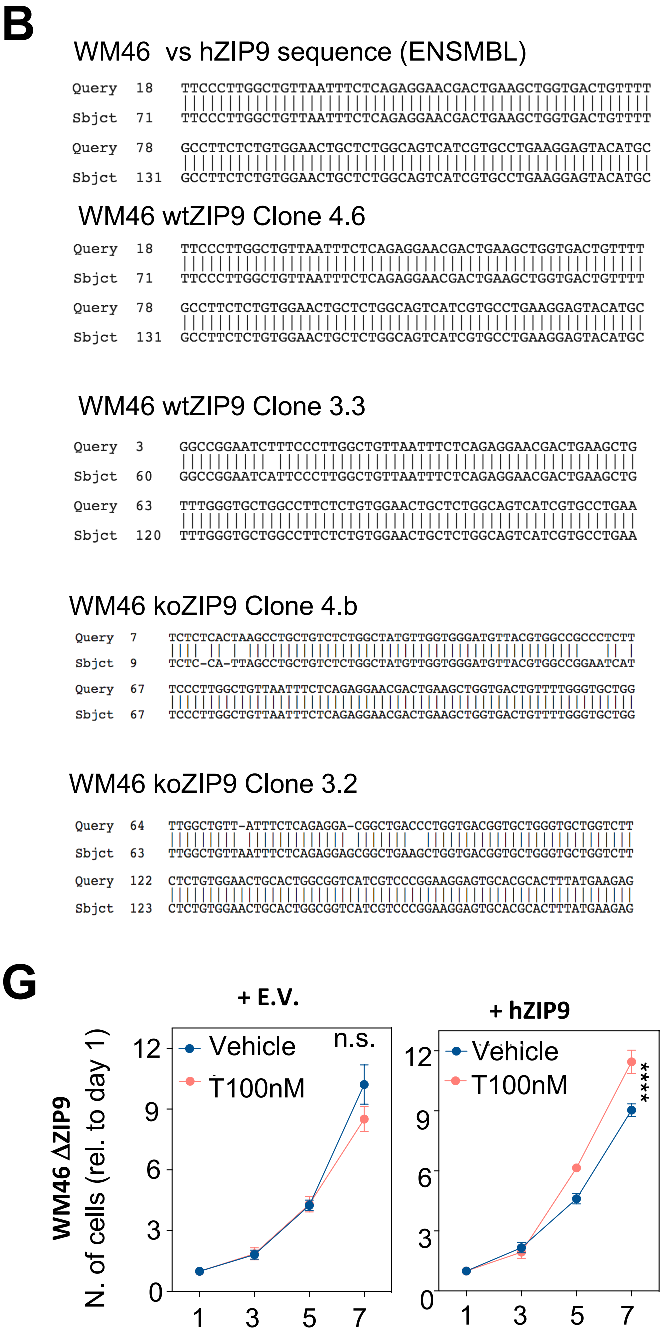


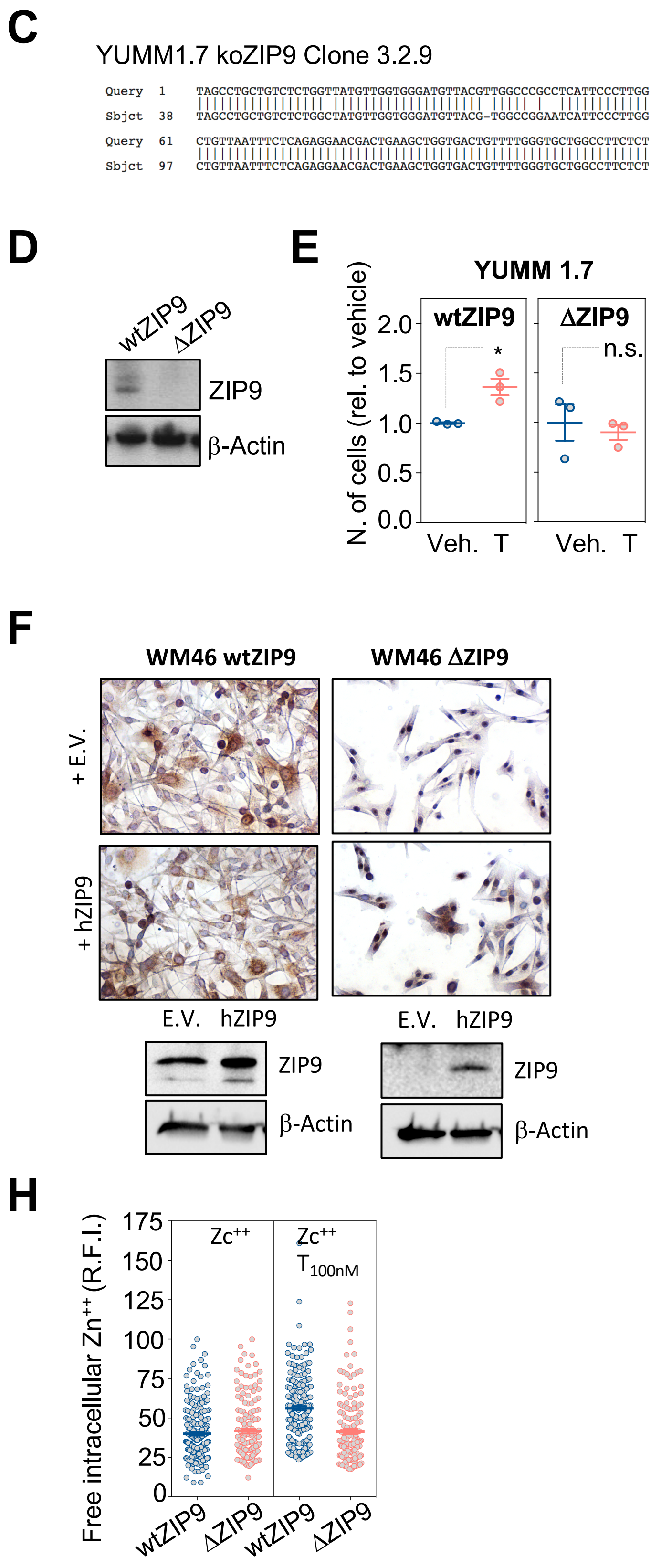
