## Supplementary Figure 7 for "ZIP9 is a Druggable Determinant of Sex Differences in Melanoma"

## A

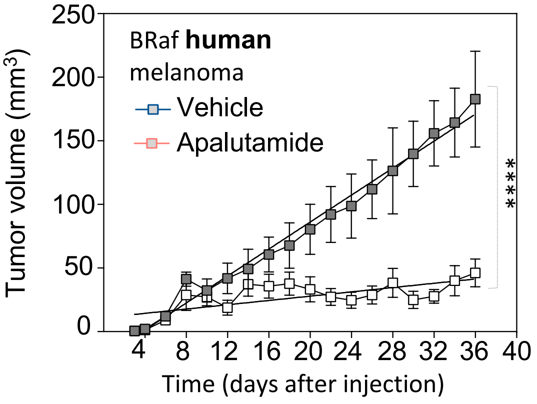


| Exponential growth equation **Best-fit values** | Males | Females |
|---|---|---|
| Y0 | 20.64 | 15.5 |
| k | 0.06241 | 0.02748 |
| Tau | 16.02 | 36.39 |
| Doubling Time | 11.11 | 25.22 |
| **Std. Error** | | |
| Y0 | 4.163 | 3.06 |
| k | 0.00669 | 0.0075 |

## B

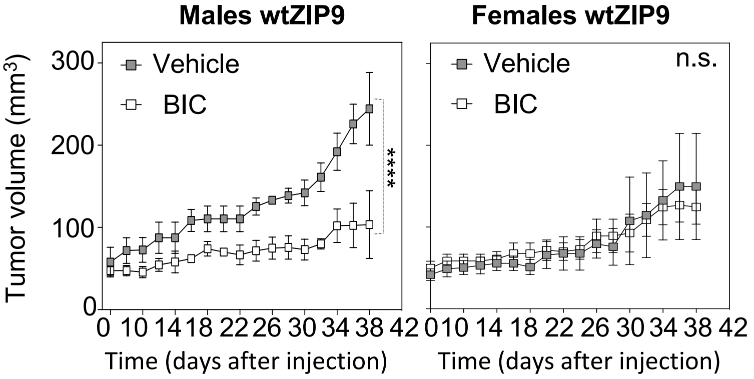


| | Vehicle | BIC |
|---|---|---|
| Y0 | 41.04 | 38.86 |
| k | 0.04079 | 0.02272 |
| Tau | 24.52 | 44.02 |
| Doubling Time | 16.99 | 30.51 |
| **Std. Error** | | |
| Y0 | 4.767 | 5.274 |
| k | 0.003447 | 0.00439 |

| | Vehicle | BIC |
|---|---|---|
| Y0 | 32.77 | 29.35 |
| k | 0.03193 | 0.03146 |
| Tau | 31.32 | 31.79 |
| Doubling Time | 21.71 | 22.03 |
| **Std. Error** | | |
| Y0 | 5.011 | 5.564 |
| k | 0.006338 | 0.00787 |

## C

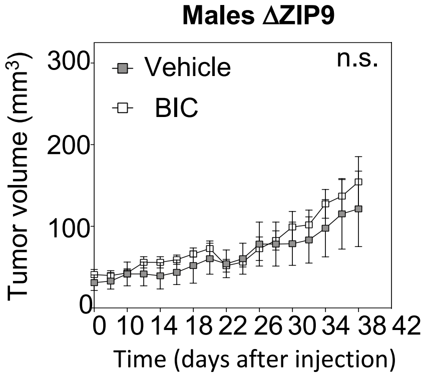


| | Vehicle | BIC |
|---|---|---|
| Y0 | 33.3 | 26.63 |
| k | 0.03102 | 0.03684 |
| Tau | 32.24 | 27.14 |
| Doubling Time | 22.34 | 18.81 |
| **Std. Error** | | |
| Y0 | 5.106 | 6.697 |
| k | 0.005979 | 0.00958 |
