## Supplementary Figure 8 for "ZIP9 is a Druggable Determinant of Sex Differences in Melanoma"

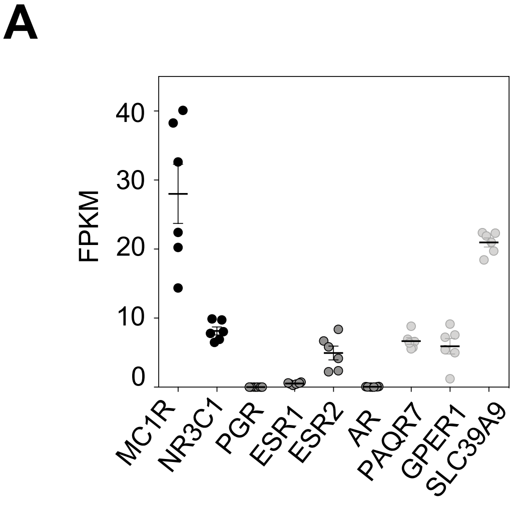


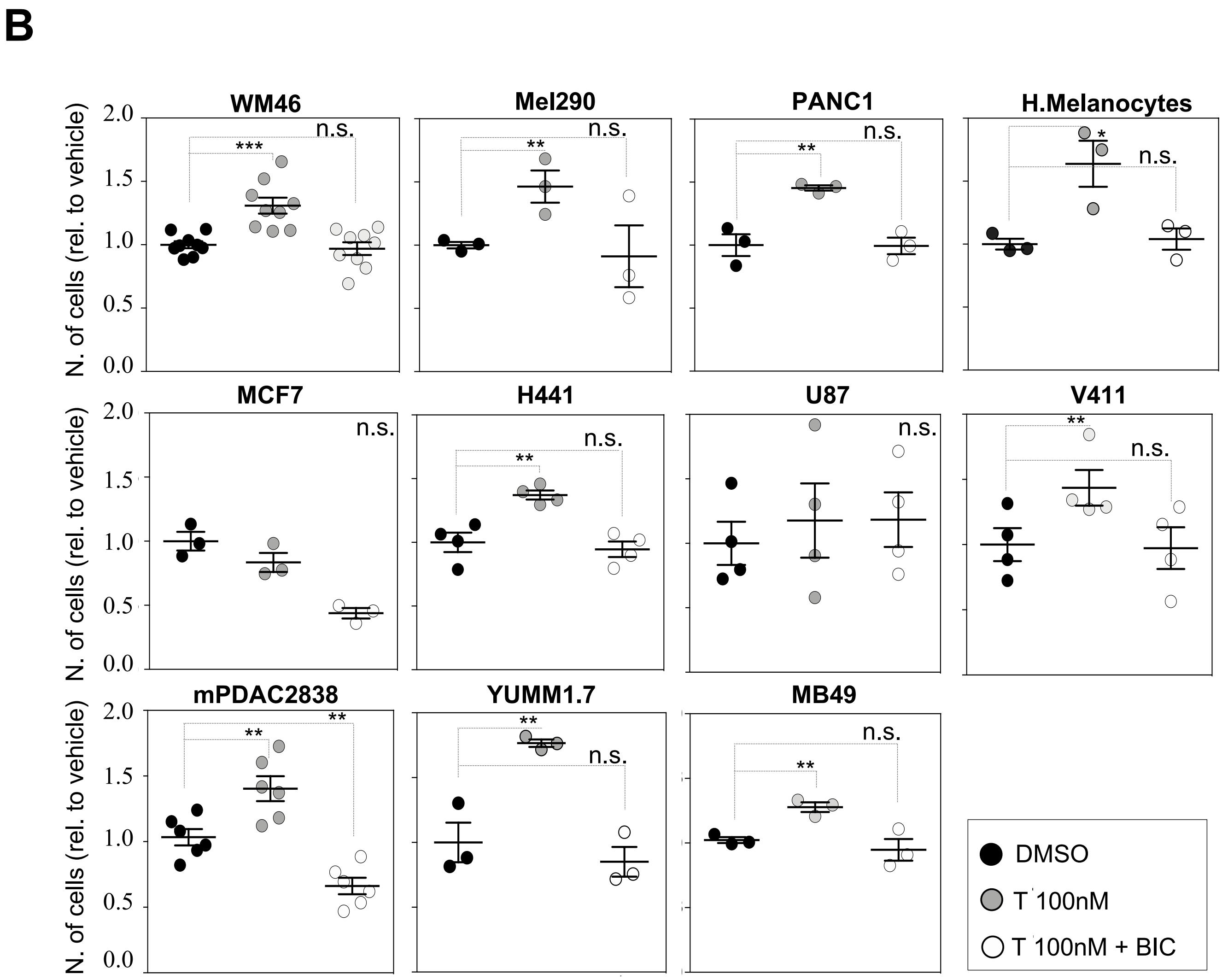


**Cell types**

**WM46:** Human skin melanoma
**Mel290:** Human uveal melanoma
**YUMMS1.7:** Murine skin melanoma
**PANC1:** Human pancreatic adenocarcionma
**mPDAC:** Murine pancreatic adenocarcinoma

**MCF7:** Human breast cancer
**MB49:** Mouse bladder cancer
**H441:** Human lung adenocarcionma
**U87:** Human glioblastoma
**V411:** Human adult acute myeloid leukemia
