## Supplementary Table 1 for "ZIP9 is a Druggable Determinant of Sex Differences in Melanoma"

| Cell line | Specie | Sex | Genotype | Stage |
|---|---|---|---|---|
| WM46 | Human | Female | B-RAF$^{V600E}$/CDK4$^{R24C}$/PTEN$^{+/-}$ | Metastasis |
| WM2664 | Human | Female | B-RAF$^{V600D}$/PTEN$^{+/-}$ | Metastasis |
| SK-MEL-2 | Human | Male | N-RAS$^{Q61R}$ | Metastasis |
| SK-MEL-3 | Human | Female | B-RAF$^{V600E}$ | Metastasis |
| SK-MEL-24 | Human | Male | B-RAF$^{V600E}$ | Metastasis |
| SH-4 | Human | Female | B-RAF$^{V600E}$ | Metastasis |
| Yumm 1.7 | Mouse | Male | B-RAF$^{V600E/wt}$/PTEN$^{-/-}$ /CDKN2$^{-/-}$ | Primary |

**Table S1.** Features of the human derived melanoma cells used across this study. Species, sex, genotype and stage diseased are detailed for each cell line.
