## Supplementary Table 2 for "ZIP9 is a Druggable Determinant of Sex Differences in Melanoma"

| GEO ID | Platform | No. of samples | Death event | Median overall survival (months) | Ages (years) | Gender (male/female) | Primary Metastatic | Stage (I/II/III/IV) | p-value | HR | 95% CI |
|---|---|---|---|---|---|---|---|---|---|---|---|
| GSE22155 | GPL6102<br>GPL6947 | 70 | 60 | 7.27 (2.10–13.80) | 56.63 ± 14.58 | 39/31 | 0/70 | 0/0/3/67 | **0.1838** | **2.32** | 0.6706-8.0268 |
| GSE98394 | GPL16791 | 51 | 18 | 93.50 (35.00–111.00) | NA | 31/20 | 51/0 | 12/22/10/0‡ | **0.0456** | **2.6873** | 1.0194-7.0843 |
| GSE19234 | GPL570 | 38 | 24 | 38.08 (23.57–65.90) | 62.66 ± 17.86 | 24/14 | 0/38 | 0/0/34/4 | **0.3953** | **0.62** | 0.2094-1.8544 |
| GSE53118 | GPL6884 | 79 | 47 | 79.74 (28.81–120.05) | 55.49 ± 15.27 | 50/29 | 0/79 | 0/0/79/0 | **0.2648** | **1.4332** | 0.7613-2.6981 |
| TCGA | Ilumina<br>HiSeqV2 | 470 | 216 | 34.45 (14.90–75.17) | 58.22 ± 15.73 | 290/180 | 103/364‡ | 77/140/171/23‡ | **0.4719** | **1.113** | 0.8314-1.49 |
| **Total** | | **1085** | **563** | **39.30 (15.92–88.00)** | **59.14 ± 15.55** | **609/394** | **221/851** | **131/215/268/149** | **0.1114** | **1.2039** | **0.958-1.5128** |

**Table S2.** Features and statistical significance of the survival analyses extracted from *OSskm*
