## Supplementary Table 4 for "ZIP9 is a Druggable Determinant of Sex Differences in Melanoma"

| | Gene | Sequence 5'-3' |
|---|---|---|
| **1** | β-Actin_Fw | AGACGCAGGATGGCATGGG |
| | β-Actin_Rv | GAGACCTTCAACACCCCAGCC |
| **2** | YAP1_Fw | CGCTCTTCAACGCCGTCA |
| | YAP1_Rv | AGTACTGGCCTGTCGGGAGT |
| **3** | LATS2_Fw | ACATTCACTGGTGGGGACTC |
| | LATS2_Rv | GTGGGAGTAGGTGCCAAAAA |
| **4** | CRM1_Fw | GCACCTCTTGGACTGAATCG |
| | CRM1_Rv | AAGCGACAGCACACACACAC |
| **5** | CDC6_Fw | AGCCTCGCATCCTATACAACC |
| | CDC6_Rv | TTCTTTCACAAGGCGGCACTC |
| **6** | CYR61_Fw | AGCCTCGCATCCTATACAACC |
| | CYR61_Rv | TTCTTTCACAAGGCGGCACTC |
| **7** | THBS1_Fw | TTGTCTTTGGAACCACAC CA |
| | THBS1_Rv | CTGGACAGCTCATCACAG |

**Table. S3:** Primers used for Real-Time Quantitative PCRs.
